# Influence of drought stress on Alfalfa yields and nutritional composition

**DOI:** 10.1101/140665

**Authors:** Yinghao Liu, Qian Wu, Gentu Ge, Guodong Han, Yushan Jia

## Abstract

It is predicted that climate change may increase the risk of local droughts, with severe consequences for agricultural practices. Here we report the influence of drought on alfalfa yields and its nutritional composition, based on artificially induced drought conditions during two field experiments. Two types of alfalfa cultivars were compared, Gold Queen and Suntory. The severity and timing of a drought period was varied, and the crop was harvested either early during flowering stage, or late at full bloom. The obtained dry mass yields of Gold Queen were higher than Suntory, and the first was also more resistant to drought. Early harvest resulted in higher yields. Decreases in yields due to water shortage were observed with both cultivars, and the fraction of crude protein (CP) decreased as a result of drought stress; this fraction was higher in Gold Queen than in Suntory and higher in early harvest compared to late harvest. Severe drought late in spring had the highest effect on CP content. The fraction of fibre, split up into neutral detergent fibre (NDF) and acid detergent fibre (ADF) increased as a result of drought and was lower in early harvested plants compared to late harvest. Suntory alfalfa produced higher fibre fractions than Gold Queen. The fraction of water-soluble carbohydrates (WSC) was least affected by drought. It was consistently higher in Gold Queen compared to Suntory alfalfa, and late harvest resulted in higher WSC content. In combination, these results suggest that the nutritive value of alfalfa will likely decrease after a period of drought. These effects can be partly overcome by choosing the Gold Queen cultivar over Suntory, by targeted irrigation, in particular in late spring, and by harvesting at an earlier time.

## INTRODUCTION

Grassland remains an important feed source for ruminant nutrition, with its high productivity and good fodder quality^1^, but alfalfa is often a necessary feed additive or alternative, especially suitable for feed production under nitrogen-limiting conditions, due to the plant’s ability to fix atmospheric N^2^. With the increase of energy costs, fertiliser (as an artificial source of soil N) has become more expensive, a trend that is expected to continue in the future, which will likely further increase the need of legume production, including alfalfa^3,4^. Agricultural forage production depends on an adequate water supply^5^, a dependence that can become problematic in semi-arid climates, especially where local effects due to climate change increase the probability of summer droughts^6–8^. Insufficient water supply can have strong effects on the production of forage legumes^9^, resulting in a decrease in yield depending on the severity and duration of drought stress^10,11^.

It is well known that forage legumes differ in drought stress sensitivity^12^. White clover is one of the most important legumes in agricultural production, but it is also relatively drought sensitive^13^. There is limited and inconsistent knowledge available about the influence of drought stress on the nutritive value of alfalfa. Under such conditions, concentrations of acid detergent fibre (ADF) and neutral detergent fibre (NDF) were found to be reduced in a range of forage legumes, while inconsistent changes were reported in crude protein (CP) concentrations^14^. A different study described an increase in ADF with a minor effect only for CP and NDF concentrations in red clover and alfalfa^15^. Another study reported an increase in water-soluble carbohydrates (WSC) under water shortage in two cultivars of soybean but it did not include alfalfa^16^. For clover species, only a small drought-induced effect on WSC was observed^17^. Alfalfa is possibly less sensitive to drought than white clover, but more research is needed to predict the influence of drought stress on the nutritive value of alfalfa, which was therefore one of the aims of this study.

Alfalfa was used here to examine the effects of drought stress on the concentrations of CP, WSC and the fibre components NDF and ADF, which were chosen as indicators for nutritive value. The CP concentration is an essential component for ruminant nutrition, and is typically high in alfalfa due to effective N fixation; WSC have a positive influence on fodder intake and are important for an efficient utilisation of dietary N; NDF content provides an estimate of the cellulose, hemicellulose and lignin content and is inversely related to voluntary fodder intake; finally ADF includes lignin and cellulose and is negative correlated with cell wall digestibility^18–2016–18^. In the present study the following questions were addressed: (1) do different alfalfa species differ in their response to drought stress; and (2) what is the effect of timing, duration, and intensity of water restriction on the nutritional parameters of alfalfa.

### Results

In two field experiments, conducted in two consecutive years, two cultivars of alfalfa were subjected to various levels of drought: Gold Queen alfalfa and Suntory alfalfa. The plants were harvested at two developmental stages. In Experiment I, conducted in 2013, two drought levels were simulated during spring (moderate drought in condition C1, and severe drought in C2) and two drought levels were applied in summer (moderate, C3, and severe drought); optimally watered plots (CO) served as control. In Experiment II, conducted a year later, severe drought was implemented either in early spring (C5) or in late spring (C6), while optimal watering (CO) was also compared to natural rainfall without further irrigation (C7). In both experiments, for all conditions, half of the plots were harvested early (samples marked ‘E’), during the initial blooming phase, and the other half late (samples marked ‘L’) at full blooming stage. The conditions tested are summarized in **Table 1**. All plots within one experiment were harvested at the same time, and the harvested alfalfa was chemically analyzed for a number of nutritional variables.

Figure 1 shows the relative available water in the soil during treatments C1-C4 (experiment I, 2013) and C5-C7 (experiment II, 2014). A plot of local precipitation and temperature during the experiments are available as **Supplementary** Figure S1. In 2013, spring was warm and dry while summer temperatures were moderate. Yearly total rainfall amounted to 224 mm with an average temperature of 16.9°C (Fig. S1). The year 2014 started with low temperatures, followed by a cool spring, with total rainfall (327 mm) and average air temperature (15.7°C) lower than in 2013. The beginning of 2014 was relatively cool, which delayed vegetation development, while May was unusually wet with over 100 mm rainfall (Fig. S1).

**Figure 1.**
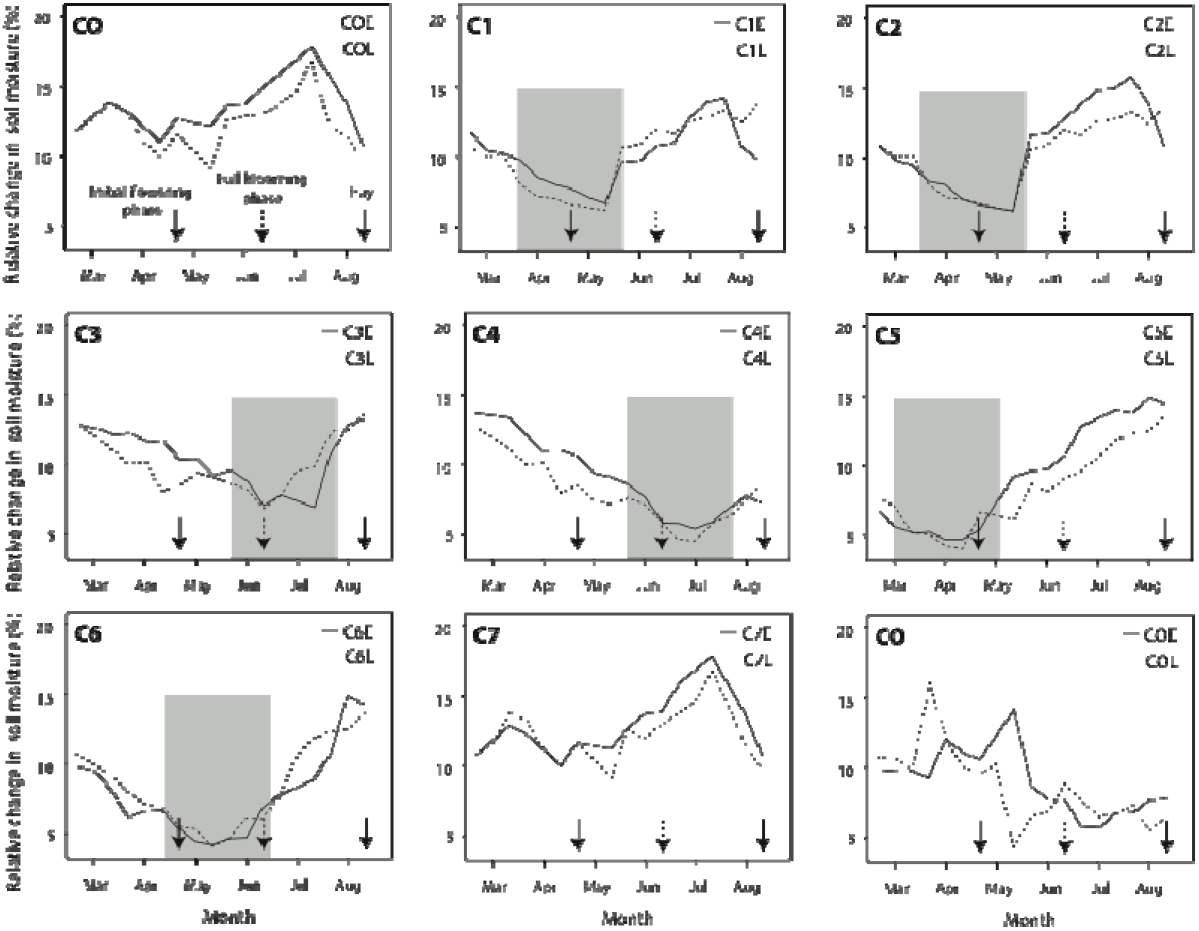
Soil moisture levels (mg water/100 mg soil) over time for optimally watered controls (CO) and experimental conditions C1-C4 (Experiment 1) and C5-C7 (Experiment 2). In each plot, two average moisture levels of four individual plots are shown, for those harvested early (solid lines) and late (dotted lines). The periods of artificially induced drought are indicated in grey. The three arrows indicate the time of early harvest, late harvest and final harvest.

The alfalfa early and late yields in Kg/ha, after removal of weeds and after drying (dry mass, DM), are shown in Table 1.

### Crude protein concentration

Crude protein concentrations from the harvested alfalfa were determined and expressed as percentage of dry mass (% DM) (**Table 2**). Whereas moderate drought applied during the spring of 2013 (C1) had no significant effect on CP concentration, severe drought (C2) resulted in significantly lower CP values when compared to optimally watered controls (CO), for both alfalfa types, and for both harvest times (Table 2). ANOVA statistical analysis indicated there was a significant difference (P<0.001) for CP content between the compared cultivars, with Gold Queen alfalfa producing significantly higher CP contents under all conditions tested. Early harvest produced significantly higher yields than late harvests, for both cultivars. The overall range of CP content of alfalfa grown with drought stress during spring varied between 15.61 and 17.61% DM. When drought stress was applied during the summer of 2013, CP content varied between 14.68 and 18.73% (Table 2), a range that was not significantly different to the range obtained with spring drought stress. The response of the two alfalfa cultivars to drought stress applied in summer was again significantly different, and CP content was lower when drought stress was severe (C4) compared to moderate stress (C3). As before, CP content was higher in Gold Queen alfalfa than in Suntory alfalfa, for all conditions tested. The ANOVA analysis further indicated that as a result of summer drought stress, Gold Queen suffered significantly more than Suntory alfalfa, resulting in a significance for DSxCV (Table 2).

In 2014, the timing of drought stress during spring was varied. A dry period early in spring (C5) reduced CP content of alfalfa compared to the optimally watered control (CO), with the exception of the Gold Queen cultivar harvested during the initial flowering phase (C5E), which actually contained a higher ratio of CP compared to the control (Table 2). A period of drought in late spring (C6) affected all plots, except for Gold Queen alfalfa harvested early (C6L). Natural rainfall without irrigation (C7) significantly reduced the CP content (Table 2). Statistical ANOVA analysis indicated that the combination of drought stress and harvest time produced significantly different results (a late harvest after late-spring drought produced lowest yields). Likewise, the combined factors of drought stress and cultivar, of harvest time and cultivar, and the combination of all three parameters were significant. The highest CP content was obtained with Gold Queen harvested early following a severe drought in early spring. The lowest CP content was obtained with Suntory alfalfa harvested late following a severe drought in late spring.

### Concentration of neutral and acid detergent fiber

The concentration of neutral detergent fibre (NDF) was higher in Suntory than in Gold Queen alfalfa, but the difference was not significant (P=0.0908). Early harvest resulted in a significantly lower fraction of NDF than late harvest (**Table 3**). Independently of the time of harvest, the content of NDF was increased by severe drought stress in spring (C2) or summer (C4), though a moderate drought during spring (C1) or summer (C3) had no effect on NDF content. During spring, both an early and a late drought (C5 and C6) increased the NDF fraction equally (Table 3). The concentration of fibre extracted with acid detergent (ADF) in part followed the same trends as those of NDF. ADF was also higher in Suntory than in Gold Queen, and this difference was now significant (Table 4). As was observed for NDF, late harvest increased the fraction of ADF, and both an early and a late period of drought during spring significantly increased the ADF fraction, as shown in Table 4. However, the ADF fraction was reduced under severe drought in spring (C2) and after a moderate drought in summer (C3), while a severe drought in summer (C4) increased the ADF fraction (**Table 4**).

### Concentration of water-soluble carbohydrates

The fraction of water-soluble carbohydrates (WSC) was significantly higher in Gold Queen than in Suntory alfalfa under all tested conditions, and late harvest produced higher fractions (**Table 5**). In Experiment I drought stress did not significantly affect these fractions, but in Experiment II, lower WSC fractions were obtained following a drought period in spring, with the exception of early harvested Suntory (Table 5).

### Hay yield reduction

At the end of the experiment all plant material was cut 3 cm above the surface and dried. Hay yields in Gold Queen alfalfa and Suntory alfalfa were reduced by all water-restricted conditions (**Table 6**), with a reduction between 27% (rain fed, C7) and 83% (Gold Queen alfalfa, draught in late spring, C6).

### Discussion

The field experiments described here were conducted to assess the effect of water restriction regimes on two alfalfa cultivars, by the use of fixed and mobile rain shelters. The validity of such an approach to study microclimatic effects has been convincingly demonstrated before^21^. We are aware that the shelters may have increased the temperature above ground, especially during hot days in summer, which may have affected plant development^22^, as it would add heat stress to the plants in addition to drought stress. Experiments conducted in winter would not suffer from this combined effect, possibly resulting in smaller changes, for instance like those reported for grain yield studied in winter wheat^23^. However, since reduced rainfall as a result of local climate changes often coincides with higher than normal temperatures, we believe our experimental conditions sufficed to investigate their combined effect. Although temporarily increased day temperatures might have negatively affected the plant growth and the yield of alfalfa, water limitation was most likely the main driver of the observed changes. Irrespective of the water supply treatment, the analytical data of the harvested alfalfa resulted in a predictive nutritive value comparable to those found in the literature^14,24^. With the values obtained, the harvested alfalfa would be considered of a moderate to high quality feed^25,26^. Under optimal water conditions, the parameters for feed quality were better for Gold Queen alfalfa than for Suntory alfalfa, with higher protein and lower fiber contents, though the higher water-soluble carbohydrate fraction in Gold Queen could be considered less beneficial.

The fraction of crude protein was reduced as a result of drought, with a similar decrease in both cultivars, but even so, CP content remained higher in Gold Queen than in Suntory. The CP fraction generally depends on the amount of available N^1,27^ and alfalfa is particularly effective in N-fixation. When the N fixation performance of alfalfa was determined, it produced 10 to 30% higher fixation levels than other legumes^28^. Thus, the degree of nitrogen fixation determines the availability of N for protein production, but it is not the only limiting factor for biomass production: obviously, this is also determined by water availability. Experiments with soybean identified N uptake as an important factor for biomass production under drought^16^, and peanut plants decreased their N fixation under drought stress^29^. We interpret the decrease in CP fractions in drought-stressed alfalfa to be caused by a combined stress response to water limitation in addition to a decrease in N fixation.

The content of neutral fiber increased under strong drought stress, a change that was also observed for acid-extracted fiber under certain conditions, though mixed results were obtained for the latter. Fiber concentration is influenced by many interacting factors, such the phase of plant development, leaf-stem ratio, environmental conditions (water, temperature, available light etc.), and availability of nutrients^24–26^. The increase in NDF and ADF fractions under stress is not supported by findings in the literature, where a delayed maturity under drought was reported, associated with lower NDF and ADF concentrations^14,25^. The major difference between the NDF and the ADF fraction is that the former included hemicellulose (the other main components are cellulose and lignin for both fractions), and the stronger and more consistent increase of the NDF as a result of drought stress in alfalfa suggests that production of hemicellulose is most affected by water restriction. However, results on the effects of drought on hernicellulose concentrations are inconsistent in the literature, as some authors have reported decreased hemicellulose concentration under drought, while other reported an increase^30^. We found that the ADF concentration was consistently lower than that of NDF, a finding that has been reported for other legumes and for most grasses as well^26^. A lower fiber concentration is generally considered beneficial, as it may lead to a higher herbage intake and to an increase in digestibility of forage. An early harvest resulted in lower fiber content and this, combined with a higher protein content, suggests that harvesting early in the season may improve the quality of the alfalfa, particularly after drought.

The fraction of WSC was least affected by drought stress, showing only a minor decrease as a result of drought stress in spring, although others have reported an increase as a result of drought in other plant species^16,31^. Gold Queen alfalfa contained significantly higher fractions of WSC, which might explain why it was also generally more capable to cope with drought stress. A high WSC concentration in plants would result in a higher osmotic potential, which drives the uptake of soil water and is therefore of importance to minimize drought stress effects^32^. This osmotic adjustment is a physiological mechanism in response to drought^16^, but in our experiments the WSC content changed marginally, only producing a significant decrease during spring drought.

Without irrigation, yields were low and nutritional parameters poor, as demonstrated with the samples grown under natural rainfall. When water supply is limited and continuous irrigation may not be possible, the timing of irrigation needs to be carefully considered. Our results indicate that the most beneficial effect can be expected if irrigation prevents a severe drought in late spring.

Digestibility of fodder may decrease under strong drought stress due to a tendency to lower WSC and higher fiber fractions, and combined with lower protein content this would redue the nitritive value. However, the decreased protein to fiber ratio in alfalfa following a drought would result in a decrease of nitrogen secretion in the urine of ruminants^20^, which can be considered beneficial for the environment. The choice of cultivar (Gold Queen) and an early harvest can minimize drought effects. Animal experiments need to be performed to further assess the feed-to-weight conversion and waste production of alfalfa grown under drought stress.

### Conclusions

The production of alfalfa is a main agricultural activity in areas of China where relatively sandy and infertile soils do not allow many other crops to be produced, but in particular these areas are expected to suffer from increased periods of drought as a result of climate change. It is therefore important to anticipate possible changes in the nutritive value of alfalfa as a result of drought stress. We have demonstrated that only severe drought stress has an impact on yield and composition of alfalfa. Strong drought led to a decrease in hay yield, a decrease in CP content, and an increase in fibre. These effects might in combination decrease the digestibility of the herbage. However, as the ratio of CP to WSC decreased under drought, this could reduce the N surplus in ruminates. We observed differences between the two tested alfalfa cultivars, both in their performance under optimal water supply and in their response to drought stress, with Gold Queen performing better than the Suntory cultivar. Finally, an early harvest could minimize the effects of drought. The reported findings may assist farmers in choosing the best cultivar, irrigation strategy and harvesting time, to mitigate the effect of decreased precipitation that can be expected in the future.

### Material and methods

#### Field experiments

A field study was conducted in 2012–2014 in a vegetation area of Ar Horqin Banner near the Nei Monggol Autonomous Region, China (coordinates 37°43’N; 120°22’E), using a randomized complete blocks design with three variable parameters tested with four replications. Two types of alfalfa were compared: Gold Queen and Suntory alfalfa. Conditions resembling severe draught and moderate draught were compared with optimal water supply whereby the timing of water restriction was also varied. The plants were harvested early (samples designated ‘E’) during the initial flowering phase, or late (‘L’) during full bloom. Details about the applied water regimes are described in **Table 1**. Drought stress was imposed during three periods with varying severity. The trials were divided into two experiments. In Experiment I (2013, spring through summer, sowing date 23 July 2012, harvest date 14 September 2013), three restricted water supply regimes were compared to optimal water supply: severe drought stress during spring, and moderate and severe drought stress during summer. Moderate stress corresponded with 20–40% usable water capacity of the soil and severe stress corresponded to only 10–15%. In Experiment II, conducted a year later (sowing date 16 August 2013, harvest date 21 September 2014), two water restriction conditions (15% available water), in early spring and in late spring, were compared to rain fed and optimally watered plots.

Drought stress was implemented by restricting rain precipitation on individual plots, using a 12m long, 6m wide, and 5m high foil cover (CASADO, Dou-ville, France). This stationary shelter was covered by 200-μm polythene foil, which was mounted over the plot. In order to attain good ventilation and to minimize microclimatic effects of the shelter, the front and the sides were left open.

#### Alfalfa types

In both experiments two alfalfa genotypes were examined; Gold Queen alfalfa is a novel American cultivar, marketed as being salt-tolerant, and suitable for high saline-alkali soil. Suntory alfalfa is a French cultivar, which has a high yield of high quality and is known to be disease-resistant.

The seeds were kindly provided by the College of Agriculture and Animal Husbandry, ChiFeng, China. The seeding density was 0.5 kg seed per 0.667 hm^2^ and plot sizes ranged from 6.6 to 8.2 m^2^.

#### Climate conditions

Air temperature and precipitation were recorded at 2 m height with a iMETOS weather station (Pessl Instruments, Weiz, Austria) located on the experimental field. The agrometeorological advisory system from the China Weather Service (CWS, 2014) was used to plan irrigation scheduling.

#### Soil composition and water content

The soil was characterized as Haplic Luvisol with an available water capacity of 120 mm (0–90 cm), and a groundwater level 10 m below surface. The soil was composed of 36% corn soil, 27% sand, 12% chernozem and 5% of other components. The pH of the soil (in CaCl_2_ suspension) measured in summer 2013 was 7.3. The soil moisture was recorded during the experiments using a portable soil moisture probe Diviner 2000 (Santé Technologies, Stepney, Australia). Plastic tubes with a diameter of 5 cm were installed to a depth of up to 150 cm. Soil moisture readings were taken at 10 cm intervals from 5 to 125 cm three times per week from the beginning of vegetation to harvest. The soil water content was also determined gravimetrically on several occasions in order to obtain a site-specific calibration (R^2^ = 0.64). The soil moisture data over time are presented as ml/100 g soil (%).

#### Sampling and measurements

For each experiment, plots were harvested by hand. For harvests, over an area of 0.09 m^2^ (in Experiment 1, 0.18 m^2^ in Experiment 2) per plot the plants were cut at a height of 3–4 cm above the soil surface and cuttings were separated from weeds immediately after harvest. Dry weight of alfalfa was determined after drying at 60°C for 72 h in a drying oven (ULM 800, Member GmbH, Schwa Bach, Germany).

For analysis of CP, NDF, ADF and WSC, dried samples were ground to 1 mm and these were analysed by near-infrared reflectance spectroscopy (NITS). All findings are reported as % dry mass (%DM). The spectra were analysed using the large dataset of calibration samples from different kinds of grasslands by the Institute VDLUFA Qualitätssicherung NIRS GmbH, Kassel, Germany.

#### Statistical analyses

Analyses of variance were carried out with the GLIMMIX procedure of SAS 9.3 (SAS Institute, Cary, NC, USA). We did a three factorial analysis of variance (ANOVA) for CP, NDF, ADF and WSC concentrations of two species in initial flowering phase and full-bloom phase of the harvest following each stress period (Payne, 2002). Experiment I and II as well as individual years were analyzed separately because of different water regimes among the years. Correlations were calculated with the CORR procedure of SAS. Graphs were created with SigmaPlot 12 (Systat Software Inc., Chicago, IL, USA). The three factors were legume species (LS), flowering phase (FS) and drought stress (DS). Relationships between selected variables were examined with a linear regression model.

## Acknowledgements

We are most grateful to all community members in the study region for their time and their confidence in our research. We also sincerely thank the staff of the provincial health department in Chifeng. This work was funded by the NSFC (Natural Science Foundation of China) under the number 31360585. The support of Zhu GD, Wang Mingchao, Fan Wenqiang, and Cheng Qiming for their help with the field experiments is gratefully acknowledged.

## Author Contributions

Liu YH and Jia YS conceived the study. Ge GT and Liu YH designed the experiments. Fan WQ, Wang ZJ, Han GD and Liu TY performed the fieldwork. Wang W and Liu XB supervised the fieldwork. Yin Q performed the quantitative data analysis. Liu YH wrote the manuscript. All authors reviewed and approved the manuscript.

## Additional Information

Competing financial interests: The authors declare no competing financial interests.

---

**Fig. S1.**
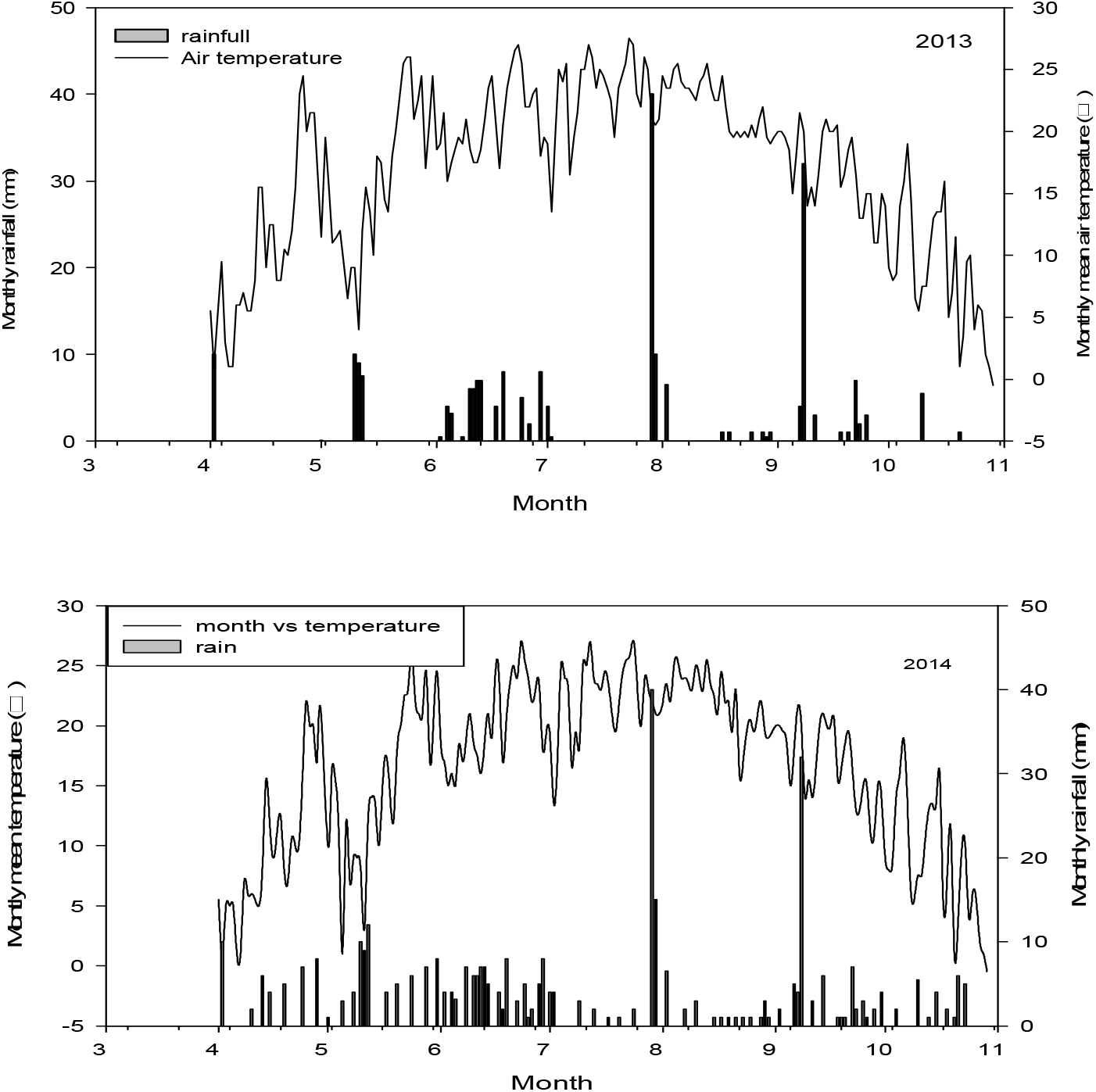
Air temperature and rainfall in the two years during which the experiments were conducted.

